# SpaMTP: Integrative Statistical Analysis and Visualisation of Spatial Metabolomics and Transcriptomics data

**DOI:** 10.1101/2024.10.31.621429

**Authors:** Andrew Causer, Tianyao Lu, Christopher Fitzgerald, Andrew Newman, Hani Vu, Xiao Tan, Tuan Vo, Cedric Cui, Vinod K. Narayana, James R. Whittle, Sarah A. Best, Saskia Freytag, Quan Nguyen

**Affiliations:** Institute of Molecular Biology, The University of Queensland, Australia; Infection and Inflammation Centre, Queensland Institute of Medical Research Berghofer, Australia; Personalised Oncology Division, WEHI, Australia; Metabolomics Australia, Bio21 Institute, University of Melbourne, Australia; School of Biomedical Sciences and Pharmacy, College of Health, Medicine and Wellbeing, University of Newcastle, Australia; Hunter Medical Research Institute, Australia; School of Biomedical Sciences, Faculty of Medicine, The University of Queensland, Australia; Department of Medical Biology, University of Melbourne, Australia; Department of Medical Oncology, Peter MacCallum Cancer Centre, Australia

## Abstract

The ability to spatially measure multi-modal data provides an unprecedented opportunity to comprehensively explore molecular regulation at transcriptional, translational and metabolic levels to acquire insights on cellular activities underpinning health and disease. However, there is currently a lack of analytical tools to integrate complementary information across different spatial-omics data modalities, particularly with respect to spatial metabolomics data, which is becoming increasingly invaluable. We introduce *SpaMTP*, a versatile software that implements an end-to-end integrative analysis of spatial metabolomics and transcriptomics data. Based in *R, SpaMTP* bridges processing functionalities for metabolomics data from *Cardinal* with user-friendly cell-centric analyses implemented in Seurat. Furthermore, *SpaMTP’s* comprehensive analysis pipeline covers (1) automated mass-to-charge ratio (*m/z*) metabolite annotation; (2) a wide range of metabolite-gene based downstream statistical analyses including differential expression, pathway analysis, and correlation analysis; (3) integrative spatial-omics analysis; and (4) a suite of visualisation functions. For flexibility and interoperability, *SpaMTP* includes various functions for data import/export and object conversion, enabling seamless integration with other *R* and *Python* packages. We demonstrated the utility of *SpaMTP* to draw new biological understandings through analysing two biological system. We believe this software and implemented methods will be broadly utilised in spatial multi-omics and spatial metabolomics analyses.

## Introduction

Fuelled by the establishment of numerous Spatial-Omics technologies and recent advances in mass spectrometry imaging (MSI) techniques, spatial metabolomics (SM) has emerged as a rapidly advancing field, providing innovative and cutting-edge insights into the metabolic activity states of cells within a spatial context [1, 2]. The most common forms of MSI include secondary ion mass spectrometry (SIMS), matrix-assisted laser desorption ionisation (MALDI) and desorption electrospray ionisation (DESI), each with distinct advantages and limitations in terms spatial resolution, detectable throughput and tissue preservation [3]. In addition, technologies such as MALDI can be implemented using specific chemical compound matrices, allowing for the detection of molecules associated with different chemical classes [3, 4]. However, in comparison to current spatial transcriptomics (ST) methods, despite the range of available technologies, adoption of SM has been hindered by the lack of purpose-built software tools that enable the analysis of large data generated by SM technologies, in particular when coupled with other spatial multi-omics datasets [1, 5].

Currently, there are a handful of open-source and commercial software tools for processing and visualising MSI data [6]. Reviewed extensively by Weiskirchen et al. (2019), many of these commercial packages have highly specific functionality and require expensive licences or dependencies, such as *SCiLS Lab* [6-10]. Alternatively, open-source packages are often widely used in the analysis of data generated from high-throughput biological assays. However, only a select few software tools are currently designed for the analysis of SM data including *Cardinal, SmartGate, pySM* and *MALDIpy [9, 11-13]*. Most of these tools provide a distinct and independent analytical approach, such as *SmartGate* which utilises artificial intelligence (AI) to perform automatic peak selection and identify spatial structures in MSI data [11]. Alternatively, *pySM* is designed for automated metabolite annotation based on false-discovery rate (FDR) [12]. The most comprehensive *Python*-based package is *MALDIpy*, which is designed for the analysis of single-cell resolution SM data exported from METASPACE [13, 14]. Whilst, this package performs general pre-processing and clustering, it is limited by a reliance on METASPACE and the use of standardised downstream analysis functions implemented by the *scapy* ST pipeline [13]. Collectively, these *Python* packages were not designed for interconnection, limiting the ability to execute a complete end-to-end SM analysis pipeline.

*Cardinal* is currently the most comprehensive tool for MSI data analysis, with a range of import, processing and analysis functions, including recent AI based implementations [9]. In comparison to other spatial-omics technologies, including ST, analytical packages such as *Seurat* commonly contain functionality for various downstream analyses that include differential expression analysis, clustering and multi-modal data integration [15]. Implementation of such downstream steps in an analytic pipeline is fundamental for drawing biological insights [15]. Currently these methodologies are still incompatible with *Cardinal*, raising the need for a comprehensive and integrative analysis tool that implements these key downstream analyses.

Another major component specific to SM analytical pipelines that is absent from almost all current packages (including *Cardinal*), is the ability to annotate relative mass-to-charge ratio (*m/z*) values with informative metabolite names [9]. The ability to detect alterations in specific metabolite levels can define the metabolic activity of certain diseases or conditions, revealing insights into possible treatment strategies [16]. Due to the vast amount of data generated by various SM technologies, often thousands of *m/z* values are detected, making it difficult to identify and interpret the specific metabolite identities of each mass [16]. The online database METASPACE and python package *pySM* are useful tools for metabolite annotation, however, offer limited capabilities of downstream analysis and integration with other analytic packages [12, 14]. Furthermore, these two tools are optimised for SM data generated via high resolution instruments such as Fourier transform ion cyclotron resonance or orbitrap analysers and do not yet apply to most data obtained with time-of-flight (TOF) analysers [14].

As metabolite identification remains limited, the combination of SM with other spatial-omics modalities promises to enhance the interpretability and learnt patterns identified from SM data through integration [13, 17]. Moreover, spatial multi-omics data are valuable for systematically studying complex biological processes across different levels of molecular regulation, including DNA, RNA, protein and metabolites [18]. Hence, integrating different modalities can provide deeper insights into complex diseases and conditions, such as cancer [19, 20]. Spatial multi-omics data including SM can be achieved either by applying different technologies separately on serial sections, or applying these technologies on the same tissue section [21]. In either case, computational analysis of this data poses challenges, including cross-omics spatial registration, multimodal molecular integration and interpretation. Currently with respect to SM, no package allows for integrative analysis with other spatial-omics data.

Here, we introduce *SpaMTP* as an open-source *R*-based package for the analysis of SM and ST data. This package expands on capacity delivered in currently available tools due to its ability to directly annotate *m/z* peaks agnostic of the data generating platform, along with performing more complex downstream analyses including pathway analysis, differential peak expression analysis, pathway analysis and cross-omics molecular interpretation. In addition, by using *Seurat* as a foundation data object, *SpaMTP* inherits a range of *Seurat* methods, enabling users to easily integrate other external tools and packages into their analysis. Overall, *SpaMTP* provides users with enhanced SM analysis methods, and the new ability to seamlessly integrate SM and ST data to elevate spatial multi-omics based research.

## Methods

*SpaMTP* is designed to facilitate a data analysis framework for matched SM and ST data (**Fig. 1A**). At the core of the package is a standardised data structure that builds on the widely-used single cell analysis *Seurat* class object [22]. This ensures efficiency of data operations due to the in-built support of sparse matrices and optimisation of memory usage. Additionally, the data structure provides the ability to retain informative metadata for both spatial locations and features, including details for each *m/z* value such as metabolite annotations, active adducts and chemical formulae (**Fig. 1A**). To facilitate flexible environments, data can be loaded in various formats including imaging mass spectrometry markup language (‘.*imzML’*) files via *LoadSM*, which expands on *Cardinals* implementation, or as a pre-processed matrix using *ReadSM_mtx*. This enables data to be pre-processed using established functionality implemented by other analytic tools, such as normalisation, spectral smoothing, baseline reduction, peak binning, filtering and alignment executed in *Cardinal* [9]. *SpaMTP* also implements various data normalisation methods such as library size log normalisation and total ion current (TIC) normalisation (*NormaliseSMData*) [23]. Quality control of pre-processed data can also be visualised using a range of *SpaMTP* plotting functions to reveal the distribution of *m/z* values across the dataset.

**Figure 1:**
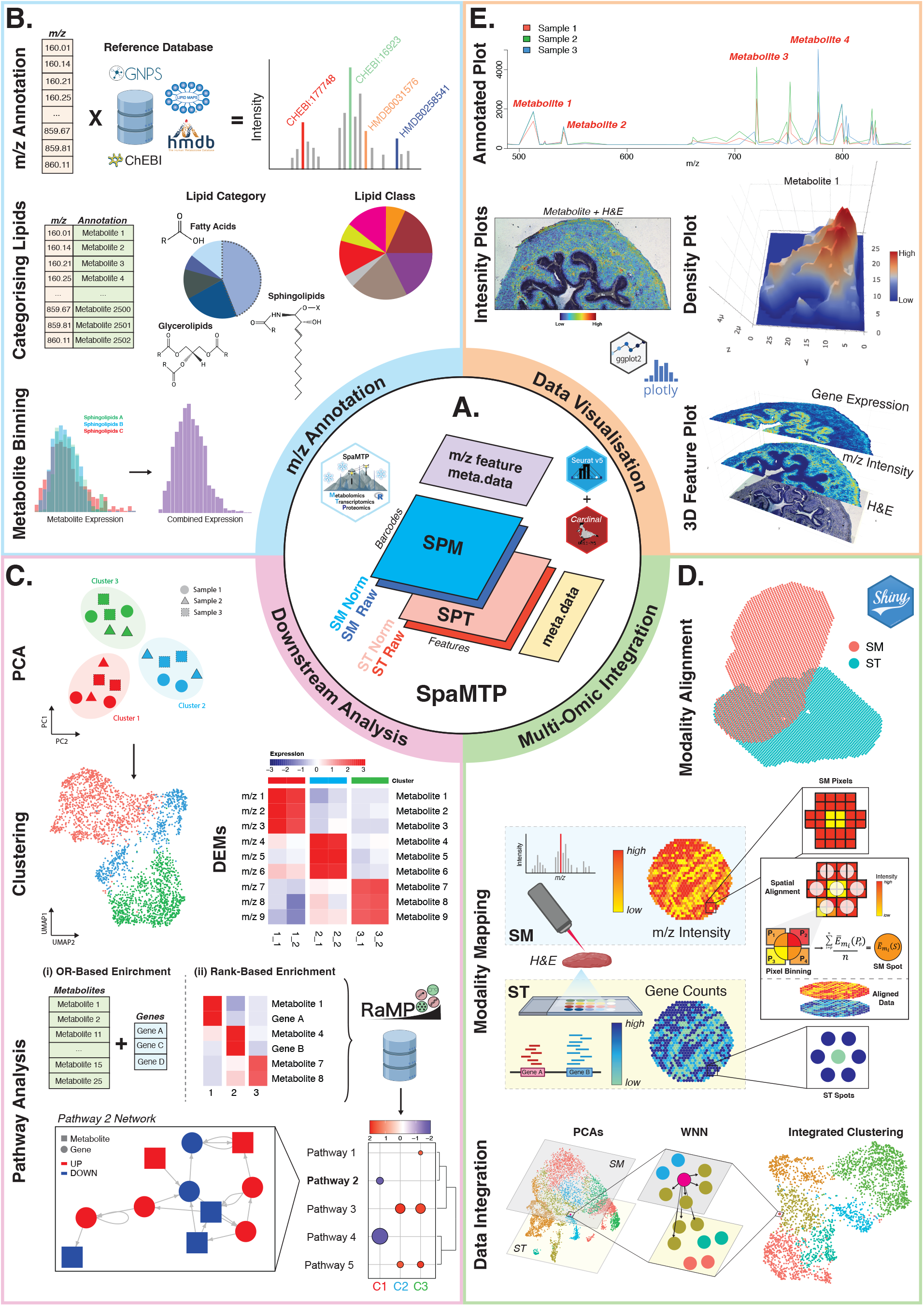
Overview of a comprehensive down-stream analytic package for spatial metabolomic and multi-omics data. **A**. Diagram of *SpaMTP’s* core object layout, capable of storing raw and processed data from various modalities. **B**. Pipeline demonstrating the automated annotation of possible *m/z* masses to relevant metabolites. Lipid annotations can be simplified using common lipid nomenclature to general lipid categories and classes. Intensity values of similar or isomer metabolites can also be binned together for plotting or analysis. **C**. Various down-stream analyses included in the *SpaMTP* package. Such as, differential metabolite expression (DEMs), PCA-based clustering and pathway analysis. **D**. Steps associated with *SpaMTP* multi-modality data analysis, including multi-omics coordinate alignment, pixel-to-spot metabolite mapping and data integration. **E**. Examples of several *SpaMTP* data visualisation methods, including intensity graphs, H&E overlayed metabolite expression figures, 3D interactive layout of *m/z* gaussian density kernel and correlated metabolite and gene expression plots.

While capable of analysing solely SM data, a novel feature of *SpaMTP* is the support of integrated multi-omics analysis. The *Seurat* ‘assay’ functionality provides the ability to store metabolic information with respect to ST data, and the corresponding H&E image aligned spatial coordinates (**Fig. 1A**). For optimal multi-modal integration, paired SM and ST data from an identical tissue section is preferred, however *SpaMTP* can also be used in conjunction with other packages to successfully align paired data from serial sections [10, 24]. *SpaMTP* inherits functionality from both *Cardinal* and *Seurat*, whilst also implementing numerous down-stream analysis methods absent from either package (**Table. S1**). These include modality alignment and mapping, integrated pathway analysis, spatial correlation analysis of both genes and metabolites, and a range of interpretable visualisation methods (**Table. S2**). As such, *SpaMTP* provides complete and flexible implementation, either free or in combination with other single-cell and spatial analysis packages.

### *m/z* annotation

A major function of *SpaMTP (AnnotateSM)* is the ability to assign putative metabolite annotations to every valid *m/z* value, applicable for any data type generated, including TOF data (**Fig. 1B**). Briefly, this is achieved through two general steps, database normalisation and database searching. First, databases were accessed as downloadable SDF files and converted to uniform text files using the R package CompoundDB [25]. The downloaded databases, including ChEBI, HMDB, GNPS and LIPIDMAPs were filtered by removing N/A and zero value entries, 47 common adducts in positive and negative mode were calculated for each neutral mass entry and appended (**Fig. 1B**) [26-29]. This resulted in a normalised internal reference metabolite database in the package suitable for annotation. Next, the reference databases are filtered by user-defined parameters such as, polarity, adducts and a filter for limiting exact mass chemical formulae by the common elements for natural products, (C, H, N, S, O, P, Br, F, Na, P, I, & Si) as defined in established methods [30]. Exact mass matching is then performed on every valid *m/z* value against the filtered reference metabolite databases by a user-defined parts per million (ppm) error. This returns a list of matches along with their corresponding information such as exact mass, formula, annotation name, InChiKey, ppm difference, and adduct type of any given match. Isomers in the reference database are summarised in terms of matching formulas and exact masses, given that a single *m/z* value may match multiple isomers from the reference databases. This representation is simplified by reporting all possible isomers per individual match against a valid observed *m/z*, providing a level of matching that corresponds to a level 4 match according to the Metabolomics Standards Initiative definition [31].

This method differs slightly to other packages, such as METASPACE which assigns annotations based on spatially informed FDR metrics [12]. Although these metrics may reduce the likelihood of false positives, unlike METASPACE, *SpaMTP* does not lose annotation preference for metabolites that do not exhibit a strong spatial pattern [12]. In addition, the user-defined threshold can be modified to decrease the chance of false positives. Each matched annotation and additional parameters are stored in the *SpaMTP m/z* metadata slot (**Fig. 1A**), providing the ability to query, label, combine and plot metabolite expression directly by name.

### Lipid classification

Although annotating *m/z* values can provide additional levels of information, limitations of different mass spectrometry equipment can result in the inability to specifically identify the metabolite based on a mass of interest. Due to the chemical structure of lipids, one *m/z* mass could represent thousands of possible metabolites [32, 33]. In addition to *m/z* annotation, *SpaMTP* builds on *rgoslin* to provide a method (*RefineLipids*) for classifying and comprehensively summarising lengthy lipid annotations into major lipid classes, using established lipid nomenclature (**Fig. 1B**). Annotated lipids can be categorised into different levels such as general lipid categories (eg. GL: Glycerolipids, SP: Sphingolipids, etc.), lipid subclasses (eg. DG: Diacylglycerols, TG: Triacylglycerols, PA: phosphatidic acid, etc.) and more refined lipid species [33]. This approach also permits intensity binning of similar lipid types, enabling the assessment of their collective biological roles rather than evaluating them individually (**Fig. 1B**).

### Interpretable clustering and spatial tissue profiling

Many spatial-omics pipelines, including *SpaMTP*, facilitate the functional interpretation of regional characteristics using principal component analysis (PCA), uniform manifold approximation projection (UMAP) and t-distributed stochastic neighbour embedding (t-SNE). *SpaMTP* implements a standard PCA pipeline (*RunMetabolicPCA*), along with providing users the ability to alter the dataset bin size to reduce noise from metabolic signal mapped to tissue. *SpaMTP* also works effectively with *Seurat* functions such as *FindNeighbors* and *FindCluters* to support feature clustering of spatial tissue bins based on metabolic signal [15]. In combination, these functions can be executed to identify spatial regions with similar metabolic landscapes and functional biological patterns, inferred from metabolic data (**Fig. 1C**).

### Pseudo-bulking differential expression analysis

Downstream statistical analyses implemented for SM data are currently limited. One key analysis method lacking is the ability to compare the abundance levels of metabolites across conditions, taking into account covariates or confounding factors. Using similar approaches to bulk and single-cell transcriptomics, here we implement a function to perform random pooling and pseudo-bulking to uncover differentially expressed metabolites (*FindAllDEMs*) [34, 35]. Based on the comparison groups specified, pixels will be randomly divided into *n* number of pools and their relative intensity values for each *m/z* will be summed (pseudo-bulking). Using *EdgeR*, pseudo-bulked intensity values for each metabolite are then compared for differential expression [35] (**Fig. 1C**). Other packages such as *Cardinal* use mean difference to identify significantly different *m/z* values, however this approach can be susceptible to dropout events and display low statistical power at small sample sizes [35, 36]. The pseudo-bulking analysis implemented in *SpaMTP* provides a more robust and scalable method of detecting differentially expressed features, by reducing noise (e.g. due to dropouts) and variance (e.g. due to random variation between individual *m/z* measures of pixels from the same sample) [37]. In addition, for each *m/z* value statistics such as fold-change and false discovery rate are calculated, permitting the use of visualisation methods such as volcano plots and heatmaps (**Fig. 1C**).

### Integrated pathway analysis of enriched metabolites and transcriptional information

*SpaMTP* incorporates two principal classes of enrichment analysis that utilise both metabolite abundance and/or gene expression; (i) over-representation based pathway analysis (ORA) (*FishersPathwayAnalysis*) and (ii) rank-based enrichment analysis (*FindRegionalPathways*) (**Fig. 1C**). These pathway approaches differ in their complexity with ORA representing the simplest and fastest method. Briefly, this function uses statistics such as Fishers’s exact test or chi-square test to determine if the proportion of differentially expressed metabolites an/or genes overlapping with a pathway’s feature set is significantly higher than the overlapping proportion of a randomly sampled feature set [38]. Rank-based enrichment analysis instead also accounts for the magnitude and direction of changes in analyte expression, allowing users to identify pathways differentially expressed between groups of interest, similar to the GSEA test [39]. In addition, *SpaMTP* performs network-based enrichment analysis (*PathwayNetworkPlots*), implementing Enrichment Map that optimises network layouts based on feature sets as nodes connected by weighted edges reflecting the biological relevance within the original pathway [40]. Here we integrate both metabolite and gene expression values to generate an interactive network visualisation of analyte interactions. This analysis is the most comprehensive, offering interpretable insights into the relative activity of given pathways, and individual analytes across a particular region.

To implement this analysis, *SpaMTP* leverages 53,952 pathway databases available through RaMP-DB, encompassing major repositories such as Kyoto Encyclopedia of Genes and Genomes (KEGG), small molecule pathway database (SMPdb), Reactome, and WikiPathways [41]. The topological structures required for network-based enrichment are derived from the omics network database provided by the Bioconductor package Graphite [42].

### Multi-modal analysis

Although *SpaMTP* can be effectively applied to solely SM data, this package is optimally designed to combine SM data with spatial data from other omics technologies. *SpaMTP* provides two functions that allows for the manual alignment (*AlignSpatialOmics*) and mapping (*MapSpatialOmics*) of spatial multi-omics data to the same common coordinate system, representing spots or cells that contain both transcriptomics and metabolic information (**Fig. 1D**).

Briefly, following the alignment of spatial coordinates to a common system, polygon objects are generated for each SM pixel. This is achieved by expanding the centroided pixel coordinate values by the median distance between neighbouring pixels. Based on the radius of each ST spot, all polygons that overlap (based on a specifiable overlapping percentage) are assigned to their respective spot. In cases where multiple pixels are assigned to a single spot/cell, the mean intensity value for each *m/z* mass is allocated. Using *Seurat’s* assay class design, the common spatial coordinate system is matched to a transcriptomics (‘SPT’) and metabolic (‘SPM’) assay, containing raw gene count and metabolite intensity values respectively (**Fig. 1A**). This results in a single *SpaMTP* object storing data values from both modalities, enabling the implementation of a range of new downstream analyses. This sample principle could be flexibly applied for integrating spatial proteomics data, forming a common object with ‘SPT’, ‘SPM’ and ‘SPP’ assays.

For example, this multi-omics object can be used to identify common genes and metabolites expressed across similar cell types. *SpaMTP* provides various functions for analysing multi-modal data including spatial correlation analysis of both genes and metabolites simultaneously. Additionally, cross-modality integration can be performed to generate informative clusters based on similar transcriptional and metabolic patterns (**Fig. 1D**). Data integration is executed using weighted nearest-neighbours implemented by *Seurat* [22], which can be performed using unsupervised or supervised (manually provided) modality weightings. This data integration can improve the clustering performance by more accurately detecting biologically meaningful cell clusters, highlighted by the multi-modal case study described below.

### Visualisation of spatial metabolic and multi-modal data

*SpaMTP* provides a large range of visualisation methods and interactive tools to intuitively display different biological results (**Fig. 1E**). To visualise pathway analysis results, *SpaMTP* provides a *ggplot2*-based dotplot to assess enrichment elements, p-values, and Jaccard distances between enriched pathways (*PlotRegionalPathways*). Additionally, *VisualisePathways* generates an expression plot alongside each pathway for ORA results, to help users identify pathways with similar expression trends and potential regions of differential expression.

For visualising the expression levels of individual mass-to-charge peaks, *SpaMTP* offers tools for summarising mass intensity and spatial intensity plots interactively. This package includes two interactive tools, *Plot3DFeatures* and *DensityPlot*, which provide a 3D visualisation (**Fig. 1E**), either displaying multi-layered metabolite/gene expression or the density of a metabolite intensity across spatial coordinates (respectively). For this, we employ in *SpaMTP* a Gaussian Kernel Density Estimation (KDE) as implemented with Javascript’s standard library with an unbiased cross-validation for bandwidth parameters [43]. These functions can help enhance the interpretability of spatially significant peaks and multi-modality based correlations.

### Implementation

To demonstrate the utility of *SpaMTP* we analysed three publicly available MALDI-TOF SM datasets. These include human lung [44], murine urinary bladder [45] and murine brain [17], where the latter contained paired 10X Visium ST data generated from the same tissue section. Full vignettes of each analysis along with full function documentation are available on the *SpaMTP* website (**Fig. S1**).

#### *SpaMTP’s* automatic metabolite annotation performance

Based on the analysis of a publicly available human lung MALDI dataset, we compared the performance of *SpaMTP’*s automated metabolite annotation protocol against previously published METASPACE results (**Fig. 2A**). All 401,325 detected *m/z* values were used to perform metabolite annotation using *SpaMTP* at a threshold of ± 3 ppm. Metabolites were named using both the HMDB [28] and LipidMaps [26] database with 80,235 and 39,215 *m/z* values being successfully annotated respectively. Comparatively, with a maximum FDR of 20%, METASPACE annotated 247 and 322 *m/z* values successfully for each respective database. Comparing overlapping *m/z* masses with corresponding annotations, approximately 90% of metabolite names matched between *SpaMTP* and METASPACE across each database (HMDB: 87%, LipidMaps: 93%). Through manual examination, at least 20% of the remaining mismatched annotations were found to be synonyms of the same metabolite (**Table. S3**). These results demonstrate the reliable performance of *SpaMTP* based *m/z* annotation, while offering convenient offline implementation.

**Figure 2:**
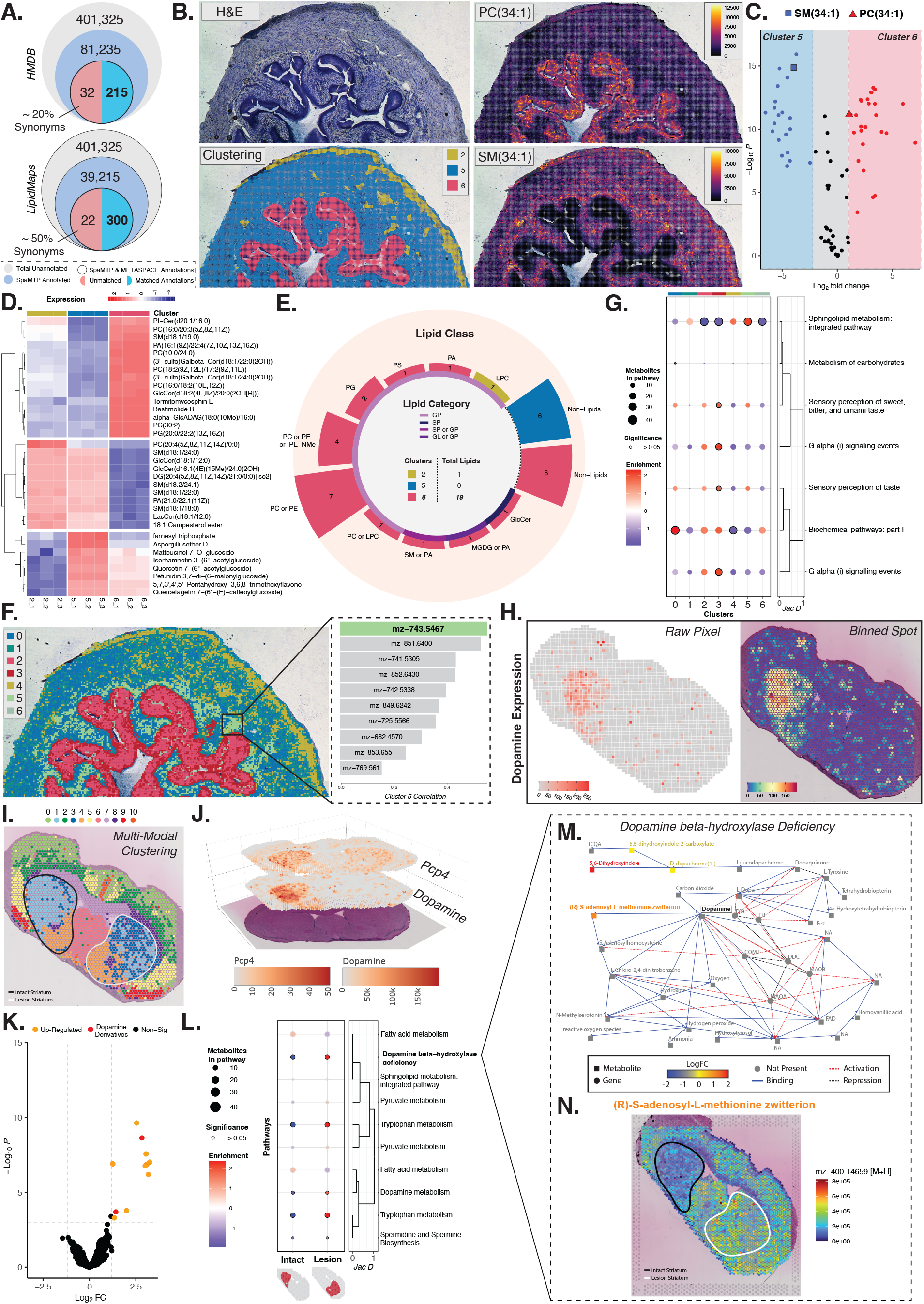
Analysis of public spatial metabolic and multi-omics datasets using *SpaMTP*. **A**. Comparative performance of *SpaMTP’s* automated metabolite annotation tool on human lung data relative to METASPACE generated annotations. **B**. Data visualisation of specific metabolite names previously identified as key structural markers of the murine urinary bladder. **C**. Volcano plot displaying differentially expressed metabolites between bladder muscle (cluster 5) and urothelium (cluster 6) regions, identified through *SpaMTP’s* pseudo-bulking analysis. **D**. Heatmap of top up-regulated metabolites for each tissue region/cluster visualised using *SpaMTP*. **E**. Circular barchart identifying high proportions of lipids expressed by the urothelium (cluster 6), grouped by lipid ‘class’ and ‘category’. **F**. Re-clustering results of the mouse urinary bladder using metabolite-based PCA embeddings estimated by *SpaMTP*, highlighting the previously unidentifiable lamina propria region (cluster 5), with top 6 most co-localised metabolites. **G**. Key significant pathways differentially expressed between re-clustered regions. **H**. Comparative spatial plots of dopamine expression on raw pixel and Visium-binned spots, from public mouse brain data with paired MALDI and 10X Visium datasets. **I**. Integrated clustering results of mapped SM and ST data utilising a weighted nearest-neighbours graph. Clustering is displayed compared to annotations, defining the relative mouse brain hemispheres containing lesioned vs intact striatum. **J**. 3D spatial visualisation of correlated gene (*Pcp4*) and metabolite (dopamine) expression across the sample. **K**. Volcano plot presenting the significantly up-regulated metabolites in the intact striatum cluster (cluster 1), which included both FMP-10 tagged dopamine molecules (yellow points). **L**. Multi-modality (transcriptomics and metabolomics) pathway analysis results representing the top significantly expressed pathways between intact and lesioned striatum regions. **M**. *SpaMTP* generated network plot depicting the ‘dopamine beta-hydroxylase deficiency’ pathway and respective components over-expressed within the lesioned striatum. **N**. Spatial expression of (R)-S-adenosyl-L-methionine zwitterion across lesioned and intact regions.

#### Murine urinary bladder down-stream analysis using *SpaMTP*

To showcase the full capabilities of a *SpaMTP*-based SM analytic pipeline, we analysed a popular benchmarking, publicly available mouse urinary bladder dataset (**Fig. 2B**) [9, 45]. Findings from the original publication identified various region-specific lipids that indicated key tissue structures of the bladder. The authors used MS/MS to confirm the exact lipids matching the respective *m/z* values. These included PC(34:1); *m/z-798*.*541*, SM(34:1); *m/z-741*.*5037* and PE(38:1); *m/z-812*.*5566*, which define the urothelium, urinary bladder adventitial layer and umbrella cells of the urothelium, respectively. Using *SpaMTP*, we analysed the pre-processed MALDI data and performed automated metabolite annotation on all detected *m/z* values. The assigned annotations for each of the above-mentioned *m/z* values correctly matched those identified in the original publication (**Table. S4**), again highlighting the efficacy of *SpaMTP’s* annotation process. These exact metabolites were plotted spatially overlaying the H&E image using *SpaMTP* (**Fig. 2B**), displaying clear tissue-specific expression patterns of lipid abundance matching the results identified in the original publication. This *SpaMTP* analysis demonstrates the ease and accuracy of identifying specific metabolites of interest, without the need for secondary experiments using more complex technologies (MS/MS).

Assessing the versatility and compatibility of *SpaMTP*, we converted the *SpaMTP* object to a *Cardinal* class object to perform spatially-aware shrunken centroid (ssc) clustering (**Fig. 2B**). This generated segments that aligned to defined tissue regions including the urinary bladder muscle (cluster 2), adventitial layer (cluster 5) and urothelium (cluster 6). A common feature of most spatial-omics pipelines to confirm clustering or segmentation results is to perform differential expression (DE) or differential abundance analysis to determine what features (metabolites/genes) differ between each group. Such a feature is not currently available in most SM-based packages. Using *SpaMTP’s* pseudo-bulking DE function (*FindAllDEMs*), we identified 70 significantly differentially expressed *m/z* peaks between our three tissue-specific clusters (**Table. S5**). Comparing between the urothelium (cluster 6) and muscle layers (cluster 5) specifically, two metabolites, PC(34:1) and SM(34:1), were found to be significantly up-regulated in these regions respectively. These two key metabolites were previously mentioned to define these specific tissue regions, supporting the accuracy of the new additional features discovered by *SpaMTP’s* DE analysis pipeline (**Fig. 2C**). To further support these findings, we can use a range of *SpaMTP* implemented visualisation tools, for example, to clearly display the over-expression of a newly identified metabolite PC(30:2) within the urothelium (**Fig. S2**).

In addition to PC(34:1) and SM(34:1), many of the DE metabolites were lipids, in particular within the urothelium region (**Fig. 2D**). Due to their molecular structure (long carbon chains), a key issue when handling lipids is the excessive numbers of possible annotated metabolites associated with one *m/z* mass (**Table. S5**). To simplify these annotations, and also determine categories and classes of lipids which are differentially expressed, we used *SpaMTP’s* lipid nomenclature functionality to refine the original annotation. These simplified annotations highlighted the large number of glycerolipids expressed within the urothelium, in particular phosphatidylcholine (PC) and Phosphatidylethanolamine (PE) lipids (**Fig. 2E**). Although there is limited research surrounding the role of specific lipids within the mouse urinary bladder, a handful of publications have identified high levels of lipids present within the urothelium of humans [46, 47].

As an alternative to ssc clustering, *SpaMTP* provides an interpretable metabolite-based PCA analysis tool. Using PCA embeddings, this sample was re-clustered using *Seurat’s* in-built graph-based Louvain clustering functions (*FindClusters*). This demonstrates a key advantage of *SpaMTP*, enabling compatibility with a plethora of established single-cell analysis methods implemented in *Seurat* to analyse SM data. Of note, the *SpaMTP-*based clustering results identified additional populations of cells which were undetectable using *Cardinal’s* ssc segmentation method (**Fig. 2F**). The urothelium region was divided into intermediate (cluster 2) and basal (cluster 3) epithelial cells, based on the spatial organisation of these two clusters [48, 49]. In addition, cluster 5 spatially matched the location of the lamina propria, a region which was acknowledged to be missing from *Cardinal’s* clustering results [9]. To effectively classify this tissue region, *SpaMTP’s* co-localisation analysis was performed (**Fig. 2F**). The top mass value that possessed spatial expression patterns correlating to the location of cluster 5 was *m/z-743*.*5467* (**Fig. 2F**). These findings support results from the original paper which defined the lamina propria based on the expression of this mass value [45]. In addition, *SpaMTP* annotated *m/z-743*.*5467* as SM(18:16), correlating with previous research demonstrating the intestinal lamina propria is associated with increased sphingomyelin levels [50].

Pathway analysis is another key method implemented in *SpaMTP*. Based on the collective differential abundance of various metabolites, pathway analysis can identify complex biological processes active within the queried sample. Focusing on the urothelium (cluster 2 and cluster 3), *SpaMTP* pathway analysis revealed G protein-coupled receptor (GPCR) alpha associated signalling pathways were highly up-regulated by both clusters relative to surrounding tissue (**Fig. 2G**). Previous research has identified various GPCRs expressed within the urothelial cells of the bladder, suggesting they are key to maintaining organ homeostasis, responding to injury and could be possible targets for treating various bladder pathologies [51-53]. Sensory perception pathways were up-regulated specifically within the basal cell layer of the urothelium (cluster 3), likely due to the sensory neurons located in the basal layer of the bladder [54]. Outside of the urothelium, regions such as the adventitia (cluster 4) and lamina propria displayed increased sphingolipid metabolism as previously mentioned [50]. The identification of these region-specific pathways demonstrates the ability of *SpaMTP* in identifying deeper biological processes, which are currently not implemented in any other SM package. Collectively these various SM-based analyses implemented by *SpaMTP* can provide users with the ability to discover novel findings about tissue biology and certain disease states, in a sensitive, robust and biologically meaningful manner.

#### Integration and Analysis of Multi-Omics Spatial Data using *SpaMTP*

While *SpaMTP* can be implemented for a purely SM based analysis pipeline, a unique feature of this package is the ability to also handle multi-modal datasets, especially for datasets with matched SM and ST assays from the same tissue section or two adjacent tissue sections. To demonstrate this functionality, we utilised a public Parkinson’s mouse brain dataset containing paired MALDI-TOF and 10X Visium data generated from the same tissue section [17]. Here, we also individually analyse two different MALDI matrices each with paired Visium data, these being 2-fluoro-1-methyl pyridinium (FMP-10) and 2,5-Dihydroxybenzoic acid (DHB) matrices respectively [17]. *SpaMTP* can handle and annotate any MALDI matrix. This includes FMP-10 data, whereby predefined metabolite annotations were assigned to corresponding *m/z* values using *SpaMTP’s AddCustomAnnotation* function. This is an essential feature, allowing users to comprehensively analyse a spectrum of metabolite and metabolic phenotypes from different chemical classes, such as neurotransmitters captured by the FMP-10 matrix and drugs and general metabolites/lipids captured by the DHB matrix [4, 17, 55]. Both Visium datasets contained ST measured per evenly distributed tissue spots of 55µm. *SpaMTP* allows for dataset alignment and mapping of metabolic signals to these spots, producing an integrated *SpaMTP* Seurat object consisting of two assays (‘SPM’ and ‘SPT’) containing SM and ST data for each individual Visium spot respectively. Due to differences in resolution between MALDI-pixels and Visium-spots, *SpaMTP* provides a function to group/bin individual pixel intensity values into a spatially colocalised spot. The spatial distribution patterns of each metabolite were unaffected by this spot-binning process, demonstrated by the comparison between spot-binned and original raw pixel expression values for dopamine (**Fig. 2H**). As a result, each spot contains both RNA and metabolite data.

The Parkinson’s model used in this sample contains an intact and lesioned hemisphere, whereby dopamine is not expressed in the lesioned striatum (**Fig. 2I**). Using ST data alone, clustering could only identify the striatum tissue region, but could not find different metabolite activity states (lesioned/intact; **Fig. S3**). Alternatively, SM FMP-10 data alone was able to correctly distinguish between lesioned states but struggled to identify common tissue structures revealed with ST data (**Fig. S3**). Both modalities were integrated with *SpaMTP* via an optimised *Seurat*-based weighted nearest-neighbours (WNN) function, that allowed for adjusting the modality weightings to prioritise metabolomics information (metabolite/gene = 0.6/0.4). This revealed two populations of spots (cluster 1 and cluster 3) which were localised within the striatum and displayed opposing metabolic activity (**Fig. 2I**). *SpaMTP’s* pseudo-bulking DE analysis of the integrated clusters revealed both FMP-10 derivatised dopamine molecules (single- and double-derivatised) were up-regulated in cluster 1 and were associated mainly with the intact striatum (**Fig. 2J**). In addition, differential gene expression analysis between all integrated clusters revealed multiple genes associated with the metabolic state of the intact striatum. These genes were unidentifiable when analysing purely ST based clustering results. *Pcp4* was the most significantly over-expressed gene within intact striatum cells, supporting results from the original publication. Pairs of metabolites and genes, such as dopamine and *Pcp4*, can be visualised in a 3D plot to clearly observe region-specific correlations between modalities (**Fig. 2K**).

Complex disease states are often regulated through diverse biological pathways involving interactions between molecular compounds across various biological modalities. In analysing the effects of striatal lesions in the mouse brain, the integration of differentially expressed metabolites and genes via *SpaMTP* revealed significant alterations in several neurological pathways (**Fig. 2L**). Specifically, using the MALDI DHB matrix and paired Visium data this study identified the elevated activity of the dopamine beta-hydroxylase deficiency pathway within the damaged striatum. Through running *SpaMTP’s* network visualisation tool (*PathwayNetworkPlots*: **Fig. S4**) to analyse the altered expression of individual genes and analytes within this pathway, a handful of metabolites were identified to play a key role in this understudied mechanism of dopamine downregulation (**Fig. 2M**). S-adenosyl-l-methionine (SAM-e), one of the pertinent metabolites linked to dopamine metabolism [56], was strongly upregulated in the lesioned hemisphere, relative to the intact striatum (**Fig. 2N**). Through the action of the enzyme catechol-O-methyltransferase (COMT), SAM-e functions as a methyl donor in the breakdown of neurotransmitters, including dopamine [56].

Since dopamine utilisation is hampered in the damaged hemisphere after striatal damage due to impaired neurotransmitter transport and reuptake, the degradation of dopamine in the synaptic cleft occurs rapidly. Excessive dopamine degradation, exacerbated by oxidative stress from high levels of reactive oxygen species induced by lesion, is likely the cause of the rise in SAM-e levels [57]. In addition, *SpaMTP’s* network analysis results also revealed another chrome indole metabolite, D-dopachrome, which is potentially formed by oxidation of dopamine when exposed to oxidative stress [58, 59]. Elevating level of D-dopachrome in the striatum is one of the markers identified in neurological disorders including Parkinson’s disease and multiple sclerosis [59]. This interplay of regulation mechanisms across different modalities highlights the potential explanation of dopamine dysregulation following striatal damage and points to the strong ability in catching crucial biological insights of the package function.

## Conclusions

*SpaMTP* is, to our knowledge, the first open-source package to implement an end-to-end analytic pipeline for integrating and analysing paired SM and ST data. Unlike other tools, *SpaMTP* provides a seamless and comprehensive implementation of numerous fundamental down-stream analyses such as data mapping, metabolite annotation, integrated clustering, differential expression and multi-modal pathway enrichment analysis. Collectively, these functionalities make *SpaMTP* a powerful tool for SM and multi-omics research providing the ability to uncover and validate novel biological insights, previously unattainable when analysing different modalities independently.

## Data Availability

All data used in this publication were publicly available. This includes the human lung MSI dataset accessible from METASPACE (https://metaspace2020.eu/dataset/2020-12-01_22h59m24s), the murine urinary bladder https://www.ebi.ac.uk/pride/archive/projects/PXD001283 and the 10X Visium and FMP-10 MALDI Parkinson’s mouse brian data https://figshare.scilifelab.se/articles/dataset/Spatial_Multimodal_analysis_SMA_-_Mass_Spectrometry_Imaging_MSI_/22770161. Databases used for metabolite annotation were downloaded from their respective locations; *HMDB* database downloaded from https://hmdb.ca/downloads (Released on: 2021-11-02), *ChEBI* database downloaded from https://ftp.ebi.ac.uk/pub/databases/chebi/SDF/ (Released on: 2023-09-01), *LIPIDMAPS* database downloaded from https://www.lipidmaps.org/databases/lmsd/download (Released on: 2023-09-15).

## Code Availability

Source code for SpaMTP is available on github https://github.com/GenomicsMachineLearning/SpaMTP. Documentation and two tutorials used to generate all figures in this publication are available through the SpaMTP website https://genomicsmachinelearning.github.io/SpaMTP/.

## Acknowledgements

Q.N., S.F., A.C. and T.L conceived the project. A.C., T.L and C.F. developed the methods and code for *SpaMTP*. A.C lead the software development. A.N., C.C., C.F., T.V., X.T. and V.K.N contributed to analysis and software development. A.C., T.L. and H.V. tested all software. A.C. curated the three public datasets, generated the vignettes, figures, supplementary material and package website. A.C. and T.L. wrote the manuscript and J.R.W., Q.N., S.A.B. and S.F. and provided feedback and revised the manuscript. Q.N. and S.F. provided project guidance and supervision. All authors read and approved the final paper.

## Ethics Declarations

The authors declare no competing interests

