## Supplementary Figures for "SpaMTP: Integrative Statistical Analysis and Visualisation of Spatial Metabolomics and Transcriptomics data"

**A.**

SpaMTP 1.0.0 | Get started | Reference | Vignettes ▾

### Getting Started

Source: `vignettes/SpaMTP.Rmd`

Listed below are a number of useful tutorials demonstrating how to use SpaMTP when analysing your spatial metabolomic datasets. For more documentation of each function used in these tutorials please visit our Reference page .

#### Useful SpaMTP Vignettes

For a general introduction in importing data, annotating metabolites and running key analysis pipelines on your spatial metabolomic data we suggest starting with the Spatial Metabolomic Analysis Tutorial. This vignette uses mouse bladder data and demonstrates how to perform general tasks such as plotting, subsetting, and manipulating `SpaMTP` Seurat objects. For those with paired multi-modal datasets we suggest working through the Multi-Modal Integration Tutorial. Here, we provide methods to align your two datasets to the same coordinates, map spatial metabolomic pixels to spatial transcriptomic spots and then run various analysis using this integrated data.

| Spatial Metabolomics Analysis using SpaMTP | Multi-Modal Integration of Spatial Data using SpaMTP | Additional SpaMTP Functionality |
|---|---|---|
| 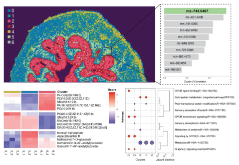 | 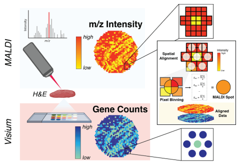 | 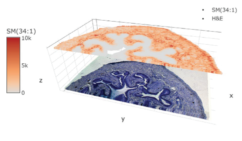 |
| An introduction to analysing spatial metabolomic datasets using SpaMTP. | Align, map and integrate spatial metabolomic and transcriptomics datasets together using SpaMTP. | Examples of additional SpaMTP visualisation and analysis functions that are useful for SM analysis pipelines. |
| GO | GO | GO |

**B.**

SpaMTP 1.0.0 | Get started | Reference | Vignettes ▾

### Reference

##### Loading Spatial Metabolic Data into a SpaMTP Seurat Object

Functions that allow the user to load in SM data in different formats

| | |
|---|---|
| loadSM() | Loads in spatial metabolomic data directly to a SpaMTP Seurat Object |
| ReadSM_mtx() | Read a Spatial Metabolomics image matrix file (.csv format) |

##### Converting Between Data Objects

Functions that allow the user to convert between SpaMTP Seurat Objects and Cardinal Objects

| | |
|---|---|
| CardinalToSeurat() | Converts a Cardinal Object into a Seurat Object |
| ConvertSeuratToCardinal() | Converts a Seurat object to a Cardinal object, including annotations and metadata |

##### Analysis of Differentially Expressed Peaks

Functions required for performing pseudo-bulking differential expression analysis

| | |
|---|---|
| FindAllDEMs() | Finds all differentially expressed m/z values/metabolites between comparison groups |
| run_pooling() | Runs pooling of a merged Seurat Dataset to generate pseudo-replicates for each sample - This function is used by run_edgeR_annotations() |

Contents


Loading Spatial Metabolic Data into a SpaMTP Seurat Object
Converting Between Data Objects
Analysis of Differentially Expressed Peaks
Visualising DEPs Analysis
Simplifying Lipid Nomenclature
Annotating m/z Masses
Metabolic and Transcriptomic Pathway Analysis
Metabolic and Transcriptomic Pathway Visualisation
Pathway Network Plotting
PCA Metabolite Analysis
SpaMTP Metabolic Data Visualisation
Additional SpaMTP Functions
SpaMTP Visualisation Helper Functions
Exporting SpaMTP Data

**Figure S1: *SpaMTP's* user-friendly website interface. A.** Example of *SpaMTP's* website containing installation instructions and multiple helpful vignettes. **B.** Sample window highlighting *SpaMTP's* detailed function documentation available on the website. The website can be found here: https://genomicsmachinelearning.github.io/SpaMTP/.

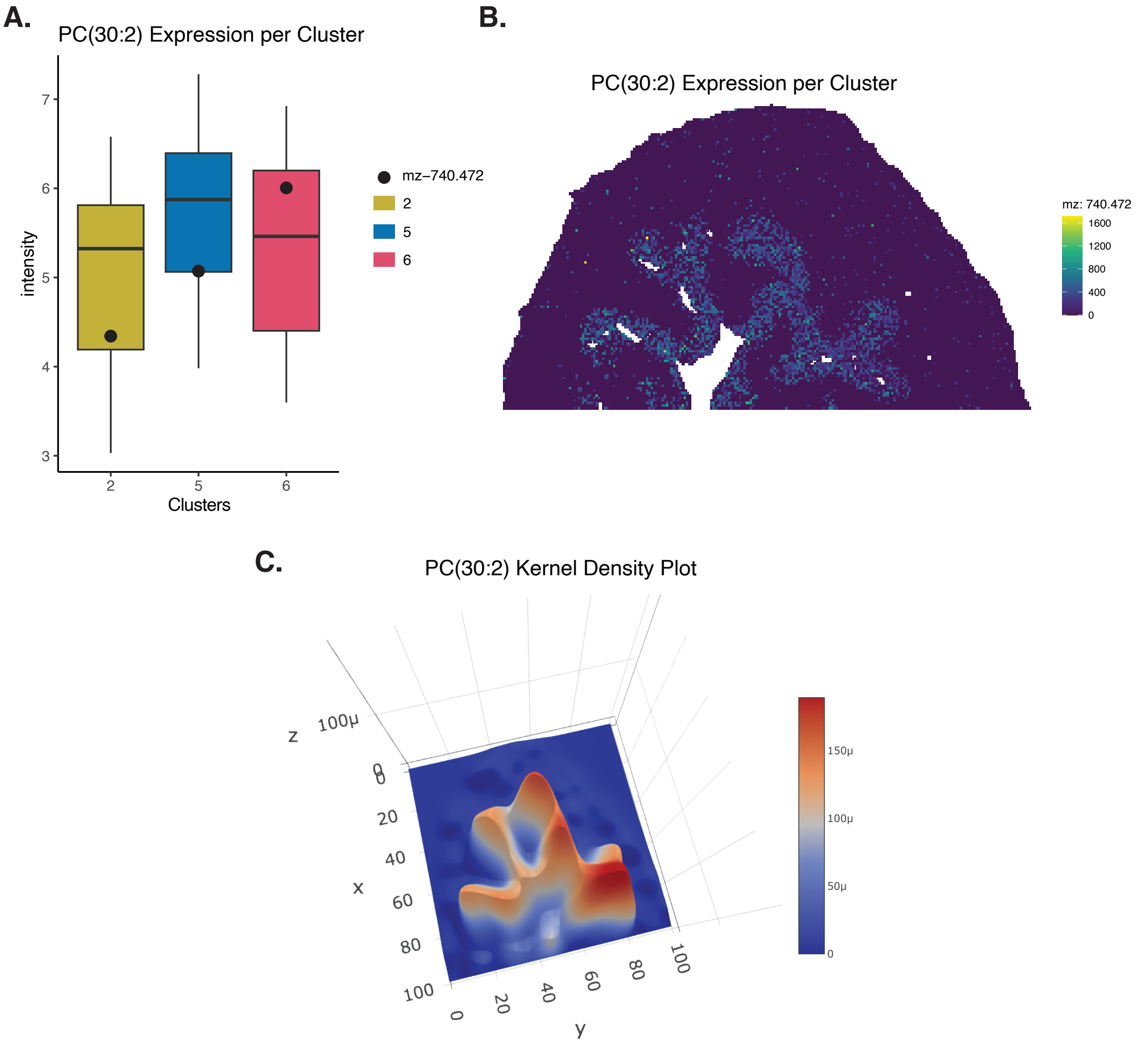


**Figure S2: Confirming PC(30:2) expression in urothelium using numerous *SpaMTP* visualisation methods. A.** Summed expression of PC(30:2) per cluster against all present *m/z* values. **B.** Spatial expression of PC(30:2) across the tissue section. **C.** Density kernel representing the gaussian distribution of PC(30:2) across the section.

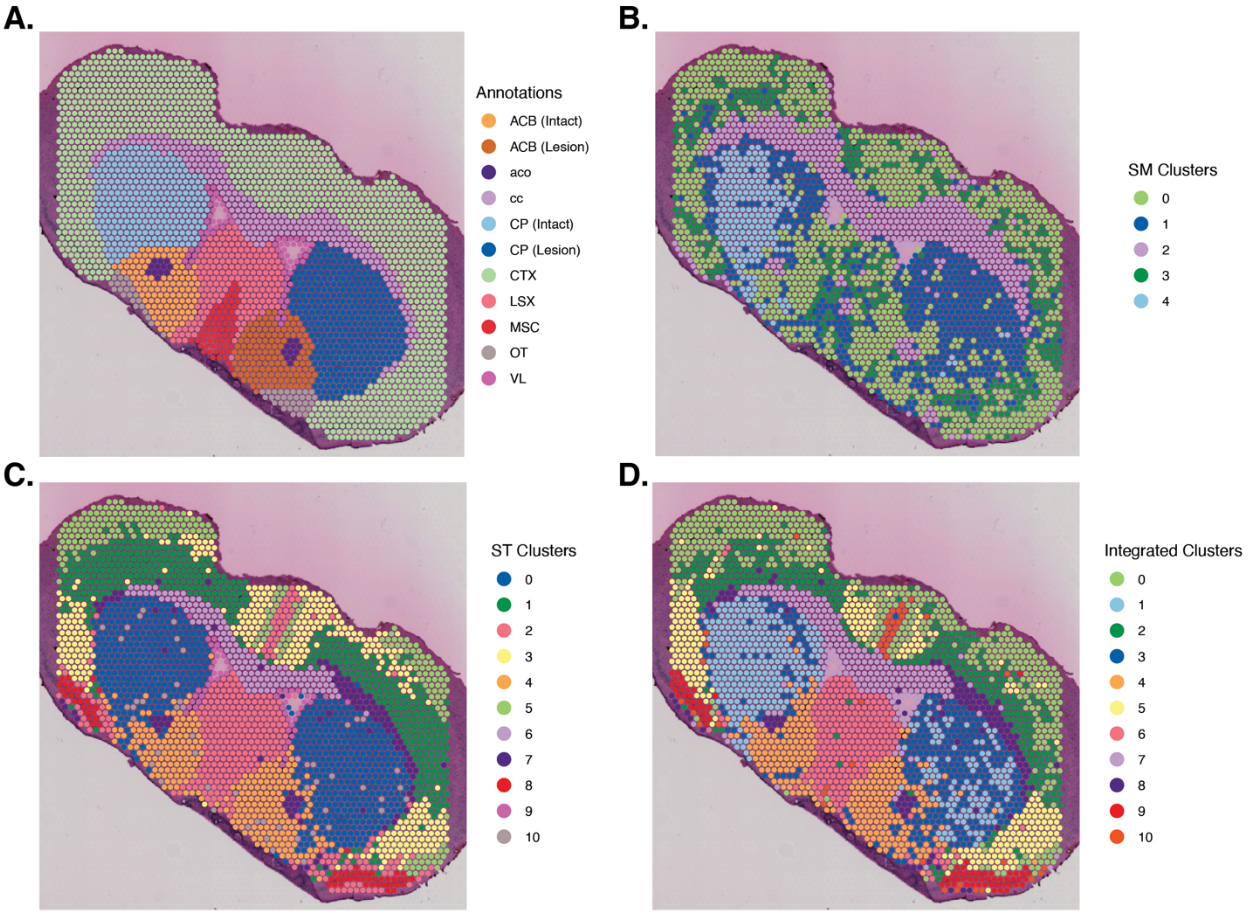


**Figure S3: Comparison of multi-omics data integration. A.** Tissue region annotations provided by the original publication. **B.** Clustering of only spatial metabolomics (SM) data. **C.** Clustering of only spatial transcriptomics (ST) data. **D.** Clustering of integrated SM and ST data using weighted nearest neighbours.

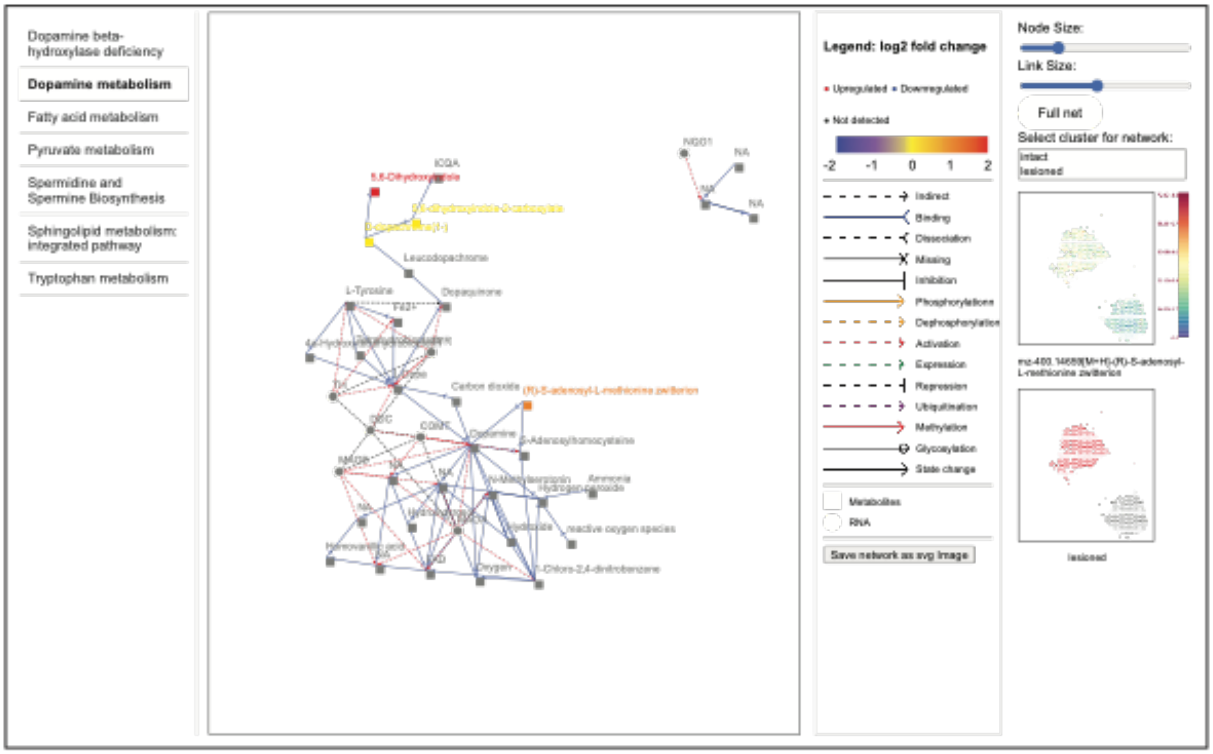


**Figure S4: Example interface of *SpaMTP's* interactive pathway network tool.** Networks of selected pathways are plotted to display interactions between analytes. Expression of individual analytes across the tissue can be visualised spatially when selected. Colours indicate the relative log fold change of each analyte with respect to the groups/clusters provided.
